# Mapping the steroid response to major trauma from injury to recovery: a prospective cohort study

**DOI:** 10.1101/577502

**Authors:** Mark A. Foster, Angela E. Taylor, Neil E. Hill, Conor Bentley, Jon Bishop, Lorna C. Gilligan, Fozia Shaheen, Julian F. Bion, Joanne L. Fallowfield, David R. Woods, Irina Bancos, Mark M. Midwinter, Janet M. Lord, Wiebke Arlt

## Abstract

**Context:** Survival rates after severe injury are improving, but complication rates and outcomes are variable.

**Objective:** This cohort study addressed the lack of longitudinal data on the steroid response to major trauma and during recovery.

**Design:** We undertook a prospective, observational cohort study from time of injury to six months post-injury at a major UK trauma centre and a military rehabilitation unit, studying patients within 24 hours of major trauma (estimated New Injury Severity Score (NISS) >15).

**Main outcome measures:** We measured adrenal and gonadal steroids in serum and 24-h urine by mass spectrometry, assessed muscle loss by ultrasound and nitrogen excretion, and recorded clinical outcomes (ventilator days, length of hospital stay, opioid use, incidence of organ dysfunction and sepsis); results were analysed by generalized mixed-effect linear models.

**Findings:** We screened 996 multiple injured adults, approached 106, and recruited 95 eligible patients; 87 survived. We analysed all male survivors <50 years not treated with steroids (N=60; median age 27 [interquartile range 24-31] years; median NISS 34 [29-44]). Urinary nitrogen excretion and muscle loss peaked after one and six weeks, respectively. Serum testosterone, dehydroepiandrosterone and dehydroepiandrosterone sulfate decreased immediately after trauma and took two, four and more than six months, respectively, to recover; opioid treatment delayed dehydroepiandrosterone recovery in a dose-dependent fashion. Androgens and precursors correlated with SOFA score and probability of sepsis.

**Conclusion:** The catabolic response to severe injury was accompanied by acute and sustained androgen suppression. Whether androgen supplementation improves health outcomes after major trauma requires further investigation.

**Précis:** A cohort study in male survivors of major trauma revealed acute and sustained androgen suppression and protein catabolism including muscle loss. Serum androgens correlated with probability of sepsis.

## INTRODUCTION

Over 5 million people worldwide die each year from serious injury (1), with almost 25% caused by road traffic collisions (RTC) (2). In England alone, there are 5400 trauma deaths and 20,000 severe injuries treated by the National Health Service annually (3). Since 2012, the establishment of 22 trauma centres in England has been accompanied by a 19% improvement in survival odds following injury (4). During this time, the UK also received severely injured military trauma patients from the conflict in Afghanistan (5, 6).

Improvements in short-term outcomes have been achieved through early resuscitation and acute care (7), often informed by approaches pioneered on the battlefield. However, improvement in survival is often offset during the weeks following acute major trauma by the systemic inflammatory response syndrome (SIRS), which is associated with increased risks of infection, multi-organ dysfunction or failure (MOD/MOF), and death (8, 9). Simultaneously, the hypothalamic-pituitary-adrenal axis (HPA) is thought to drive a hypermetabolic and overtly catabolic response. Importantly, in this profound catabolic state, patients lose valuable lean muscle and suffer from increased rates of infection and poor wound healing. Moreover, the dynamic nature of this response, especially beyond the first few days following injury and during recovery remains poorly described and understood, limiting the evidence base for novel therapeutic interventions. Burn injury also produces an extreme inflammatory and catabolic response after injury, previously targeted by anabolic steroid analogues (10) and beta-blockade (11). The dynamic changes in endogenous glucocorticoids and their influence on adrenal steroid metabolism after severe injury are not well characterized. We know that pro-inflammatory cytokines activate the enzyme 11β-hydroxysteroid dehydrogenase type (11β-HSD1) responsible for tissue-specific activation of glucocorticoids through conversion of inactive cortisone to active cortisol (12). However, only scarce data exist on what happens to early sex steroids and their precursors during this catabolic state.

To address these gaps in knowledge, we have undertaken a detailed prospective study of the endocrine and metabolic response to severe injury in military and civilian populations, recruiting patients within 24 hours of major trauma and following up for the six months post-trauma. This was undertaken to identify predictive biomarkers and therapeutic targets as well as to explore the optimal timing for therapeutic interventions that could promote better recovery after severe traumatic injury.

## MATERIALS AND METHODS

### Study Design and Protocol

This prospective cohort study was conducted in the Royal Centre for Defense Medicine and the Queen Elizabeth Hospital Birmingham, a major UK trauma centre and the primary receiving facility for UK military personnel injured abroad. Military and civilian trauma patients with an estimated New Injury Severity Score (NISS) >15 were recruited (13). NISS was used to ensure those with significant extremity trauma but lower ISS were included (14). Patients with significant head injury or pre-injury neoplastic conditions were not eligible. None of the patients received etomidate during their treatment. Informed consent was obtained from personal consultees until recovering capacity. The protocol was approved by the NRES Committee South West – Frenchay 11/SW/0177 and MOD REC 116/Gen/10.

A daily patient review allowed the injured to be clinically phenotyped. Bespoke study management software (Clinical RESearch Tool – CREST) tracked the patient and their clinical data were entered prospectively and used to calculate Acute Physiology and Chronic Health Evaluation II (APACHE II), Sequential Organ Failure Assessment (SOFA), and Simplified Acute Physiology Score (SAPS) (15, 16). Sepsis was defined using Bone’s criteria of an infection associated with SIRS (17), current at the time of the study.

Details of opioid administration were collected from the electronic health record and prescribing system (PICS) (18) on each of the study patients. The total amount of opioid given during their hospital stay was appropriately weighted and totalled with equivalence to an oral dose of 10mg morphine, adjusting for potency, delivery method and opioid preparation (19).

Blood and urine samples were collected within 24 hours of acute injury and at 3, 5, 10, 14, 21, 28 days and 2, 3, 4 and 6 months during recovery post-trauma. Blood sampling occurred between 0730 and 0900 hrs; serum was then separated and frozen at −80°C for batched analysis. We separately collected morning blood samples from 37 healthy age- and sex-matched controls, to provide a comparator for the steroid data.

### Assessment of Protein Catabolism

As a surrogate marker for muscle mass, we undertook longitudinal measurements of muscle thickness by a well validated method using portable ultrasound as described by Campbell *et al.* (20, 21). Ultrasound measurements were taken from four different muscle sites (biceps brachii, radial forearm, rectus femoris and rectus abdominis) at weekly intervals while in hospital and at 3, 4, 5 and 6 months following discharge. All ultrasound assessments were performed by two trained operators. Measurements were performed three times at each muscle site and the mean of the three measurements was recorded. The dominant arm was favoured for ultrasound assessment of muscle mass unless it was missing or unable to be measured where wounds were extensive. This non-invasive method was chosen over others such as creatine (methyl-d3) dilution (D3-creatine) (22) due to the strict dietary requirements for methy-d3 estimation, which was not practical in the context of major trauma. Similarly, we did not undertake MRI measurement of muscle mass due to the risk associated with repeated transport of critically ill patients to scanning facilities (23).

Urinary urea excretion was measured and used to estimate Total Urinary Nitrogen (TUN) excretion as described by Milner et al. (24) [Estimated Nitrogen Excretion: Urinary Urea Excretion (mmol/l) x 0.028 x 1.25 = Total Urinary Nitrogen excretion (g/l)].

### Steroid Analysis

Serum concentrations of adrenal and gonadal steroids were measured using liquid chromatography-tandem mass spectrometry (LC-MS/MS) analysis, employing a validated multi-steroid profiling method (25). In brief, serum steroids were extracted via liquid/liquid tert-butyl-methyl-ether (MTBE), evaporated, reconstituted, and analysed by LC-MS/MS for cortisol and cortisone. Serum androgens and androgen precursors (DHEA, androstenedione, testosterone) were measured following oxime derivatization (26, 27). Serum DHEA sulfate (DHEAS) was measured following protein precipitation (28, 29). Steroid metabolite excretion analysis in 24-h urine samples was carried out by gas chromatography-mass spectrometry (GC-MS) in selected-ion-monitoring (SIM) mode, as previously described (30).

Serum concentrations of sex hormone-binding globulin (SHBG) and luteinizing hormone (LH) were analysed on the Roche Modular System (Roche Diagnostics, Lewes, UK) by two-site sandwich immunoassay using electrochemiluminesence technology.

### Statistical Analysis

The raw data were evaluated by analysis of variance (ANOVA). In addition, paired and unpaired Student’s t-test, Chi-Square Analysis and Mann-Whitney tests were used where appropriate.

Generalized linear mixed-effects models (31) were used to examine the change in variables over time. Patients were included in models as random effects to account for repeat measures over time on the same individuals. Time was modelled using restricted cubic splines (32) to allow for flexible relationships (33). Severity scores were modelled as Poisson distributions due to their skewness and non-negative ranges. Plots of predicted average fixed effects with 95% confidence intervals were produced for the first four weeks and first six months post-injury as required. Analyses were conducted in R using libraries lme4, effects, rms and ggplot2.

## RESULTS

### Patient Recruitment and Clinical Characteristics of the Final Study Cohort

We screened 996 multiply injured adults. The majority of the 889 excluded patients had a NISS≤15, others had a significant head injury as their major injury component, and two were excluded due to a pre-injury diagnosis of cancer. Of the 102 patients recruited into the study, two withdrew and re-assessment in five revealed an actual NISS ≤15, leaving a study cohort of 95 patients (Fig. 1A).

**Figure 1.**
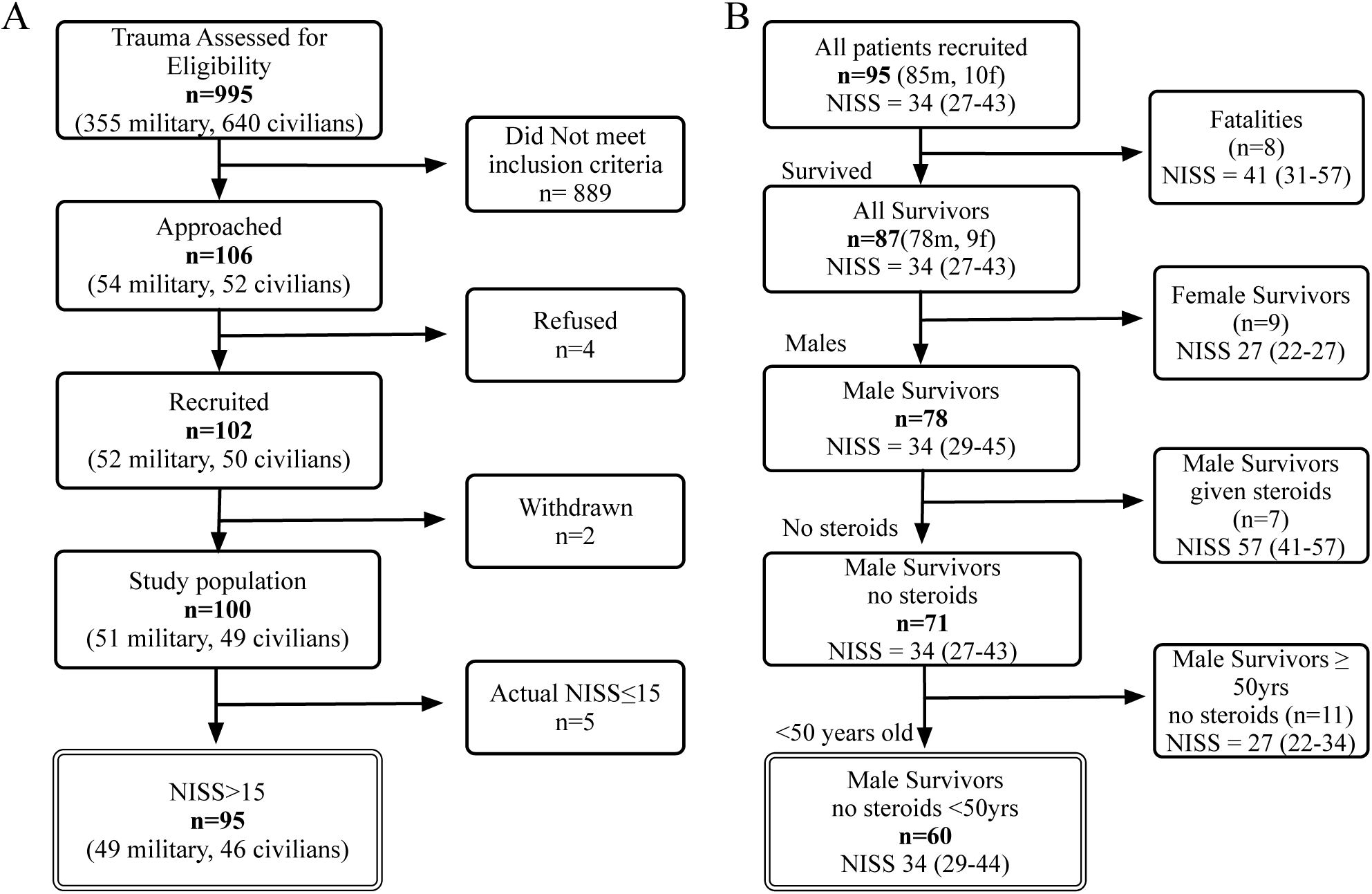
Consort diagram. (A) recruitment process and (B) subgroup selection for analysis for sixty male survivors of severe injury (NISS>15) under 50 years of age who had not been given exogenous steroids were analysed.

Excluding eight fatalities and seven patients who had received steroid therapy, 80 survivors completed sample collection over 6 months. To minimize confounders, we excluded the small groups of women (n=9) and age-advanced men (n=11), leaving our final study cohort of 60 men <50 years of age (Fig. 1B).

A summary of the cohort characteristics is shown in Fig. 2. Median age was 27 (interquartile range (IQR) 24-31) years, median NISS was 34 (IQR 29-44), and patient day-1 (=day of major trauma) APACHE II score was 21 (IQR 14-25). Patients remained ventilated on the intensive care unit (ICU) for a median of 9 (IQR 5-16) days. Median length of hospital stay was 36 (IQR 19-56) days. Improvised Explosive Device (IED) (n=33; 55%) and RTC (n=11; 18%) were the most common causes of injury. Twenty-five (42%) patients had at least one septic episode and most occurred in the second week (Fig 2C).

**Figure 2.**
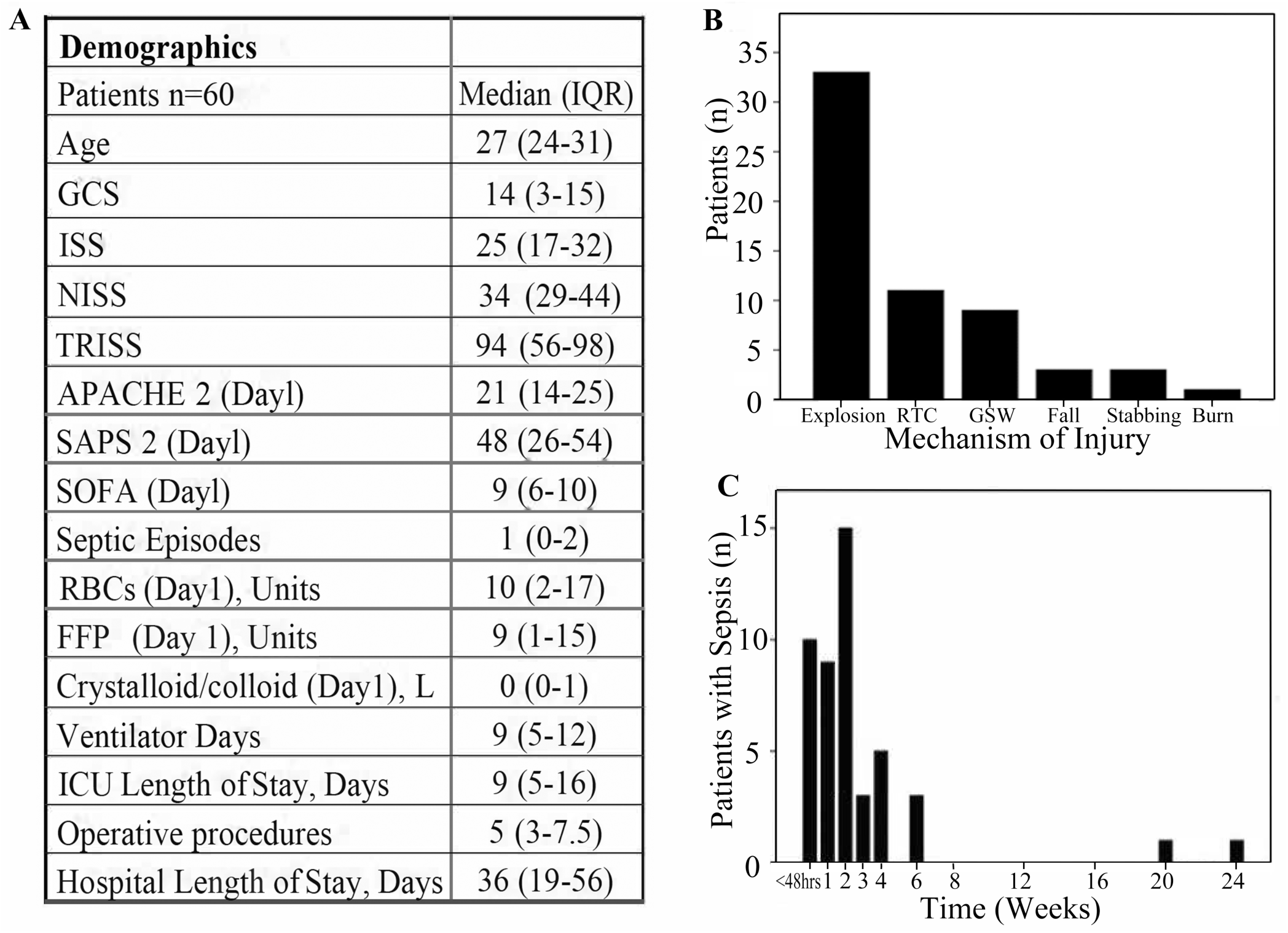
Patient Characteristics of the Analysis Cohort. (A) Demographics, (B) Mechanism of Injury and (C) the distribution of septic episodes for 60 male survivors from severe injury (NISS>15) under 50 years of age.

### Glucocorticoid Biosynthesis and Metabolism After Major Trauma

Serum cortisol concentrations increased slightly after injury, peaking at 408 (IQR 249-511) nmol/L at two weeks (Fig. 3A**; Suppl. Fig. 1A**) (34). However, concentrations remained within the wide range observed in healthy controls. Serum concentrations of the inactive glucocorticoid metabolite cortisone were lower than normal after injury, and increased slowly over time, but this trend was not significant (p=0·08) (Fig. 3B**; Suppl. Fig. 1B**) (34). The serum cortisol-to-cortisone ratio, a marker of systemic 11β-HSD activities (Fig. 3C), peaked at two weeks post-injury and returned to normal at around eight weeks. Consistent with these findings, urinary steroid metabolite excretion analysis revealed an increase in glucocorticoid metabolite excretion in weeks 2, 4 and 8 after major trauma, alongside changes in steroid metabolite ratios indicative of increased systemic 11β-HSD1 and decreased 11β-HSD2 activities, as assessed by (5*α*-tetrahydrocortisol + tetrahydrocortisol)/tetrahydrocortisone and cortisol-to-cortisone ratio, respectively (**Suppl. Fig. 2+3**) (34).

**Figure 3.**
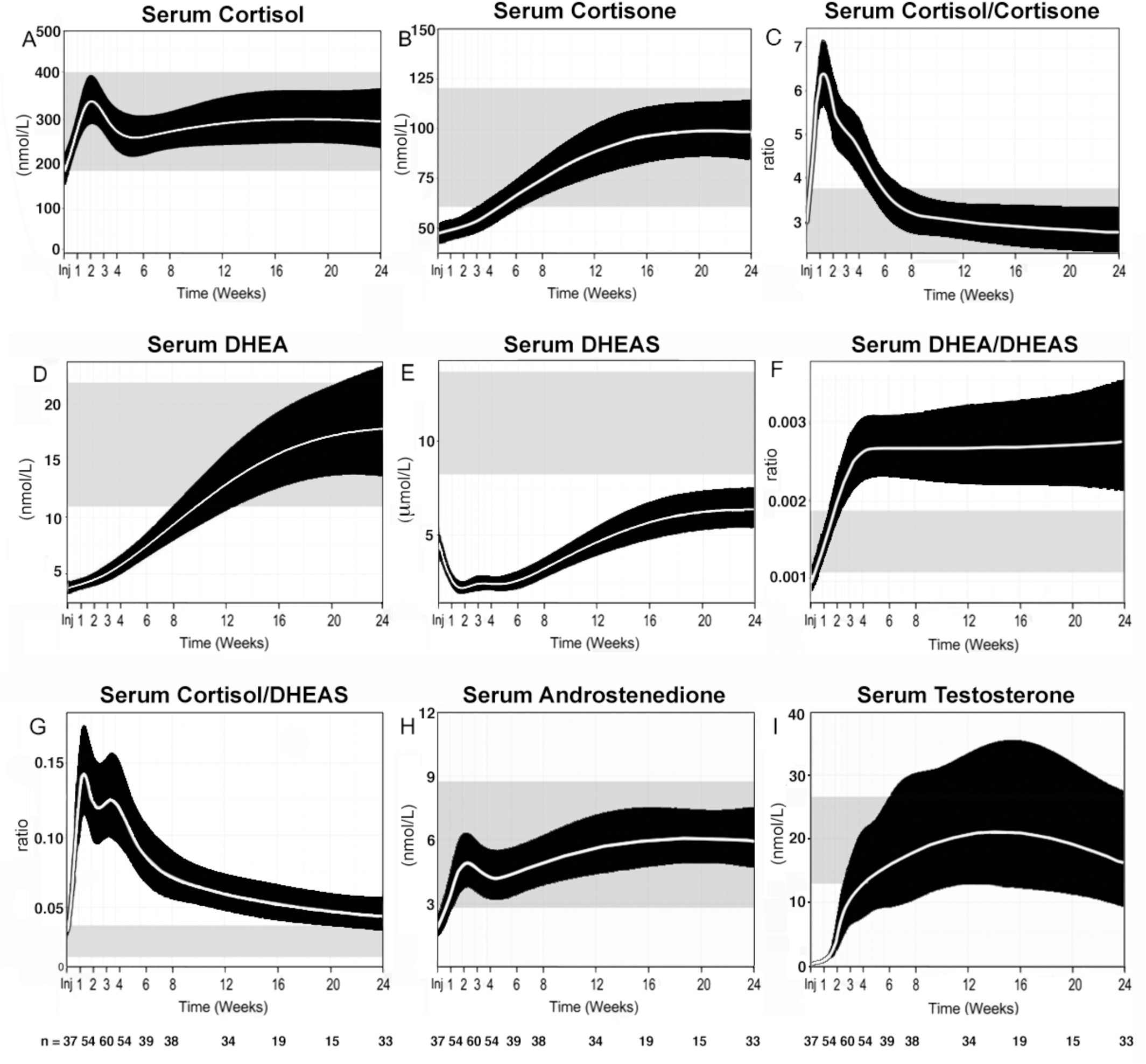
Serum steroids in 60 male survivors of severe injury (NISS>15) under 50 years of age. Serum concentrations shown include cortisol (A), cortisone (B), the cortisol-to-cortisone ratio (C), DHEA (D), DHEAS (E), the DHEA-to-DHEAS ratio (F), the cortisol-to-DHEAS ratio (G), androstenedione (H), and testosterone (I). Data are represented after modelling of the raw data (Suppl. Fig. 1) using a non-linear mixed effects model that accounts for unbalanced repeated measures using a 4-knot cubic spline. Modelled data are shown as means and 95% confidence intervals.

### Androgen Biosynthesis and Activation After Major Trauma

Serum concentrations of the adrenal androgen precursor dehydroepiandrosterone (DHEA) were very low after injury (p<0·0001, compared with healthy controls) but recovered to the normal range by three months post-injury (Fig. 3D**, Suppl. Fig. 1**) (34). In contrast, its sulfate ester, DHEAS, demonstrated sustained suppression; median serum DHEAS concentrations did not recover to values within the healthy reference range, even at the end of the 6-month study period (Fig. 3E). Consequently, the serum DHEA-to-DHEAS ratio (Fig. 3F) increased by week 2 compared with controls and failed to return to normal during the 6-month study period. The serum cortisol-to-DHEAS ratio (Fig. 3G) increased post-injury, peaking at 2 weeks, followed by a gradual decrease, but without returning to normal by the end of the 6-month study period.

Serum concentrations of the androgen precursor androstenedione (Fig. 3H) were below the reference range immediately after injury, recovering to the mid reference range at 2 weeks post-injury. Thus, serum androstenedione concentrations recovered much faster than DHEA, suggestive of rapid downstream activation of DHEA to androstenedione.

Serum testosterone (Fig. 3I**, Suppl. Fig. 4A+B**) (34) was very low following injury, starting to increase after two weeks, and recovering to the healthy sex- and age-matched reference range approximately eight weeks after injury. This was mirrored by acute suppression of serum LH immediately after injury, followed by recovery to the normal range approximately 2 weeks after injury (**Suppl. Fig. 4C+D**) (34). Serum sex hormone-binding globulin (SHBG) (**Suppl. Fig. 4E+F**) (34) concentrations were subnormal immediately post-injury, but quickly returned to the healthy reference range between injury and day 7.

Consistent with the observed decrease in circulating androgens, 24-h urinary steroid metabolite excretion analysis revealed a steep decrease in the major androgen metabolites androsterone and etiocholanolone at 2, 4 and 8 weeks (**Suppl. Fig. 4A+B**) (34). Similarly, urinary DHEA excretion, representing the sum of unconjugated DHEA and DHEA sulfate, sharply decreased to very low concentrations at 2, 4 and 8 weeks, with a transient increase in 16*α*-hydroxylation of DHEA at 2 weeks (**Suppl. Fig. 4C+D**) (34), possibly linked to the systemic decrease in DHEA sulfation (Fig. 3D-F). The overall decrease in androgen production was paralleled by a profound decrease in systemic 5*α*-reductase activity (**Suppl. Fig. 4E+F**) (34), and hence in androgen activation, as 5*α*-reductase is responsible for converting testosterone to the most potent androgen 5*α*-dihydrotestosterone.

### Protein Catabolism After Major Trauma

The 24-hour total urinary nitrogen (TUN) excretion increased immediately after trauma, peaking at 25.0±16.1 g/day at the end of the first week, returning to below 15·0 g/day by week-4. The mean maximum rate of nitrogen excretion was 33.0±21.3 g/day (Fig. 4A). The normalization of TUN excretion coincided with the gradual recovery of adrenal and gonadal androgen production (Fig. 4B+C).

**Figure 4.**
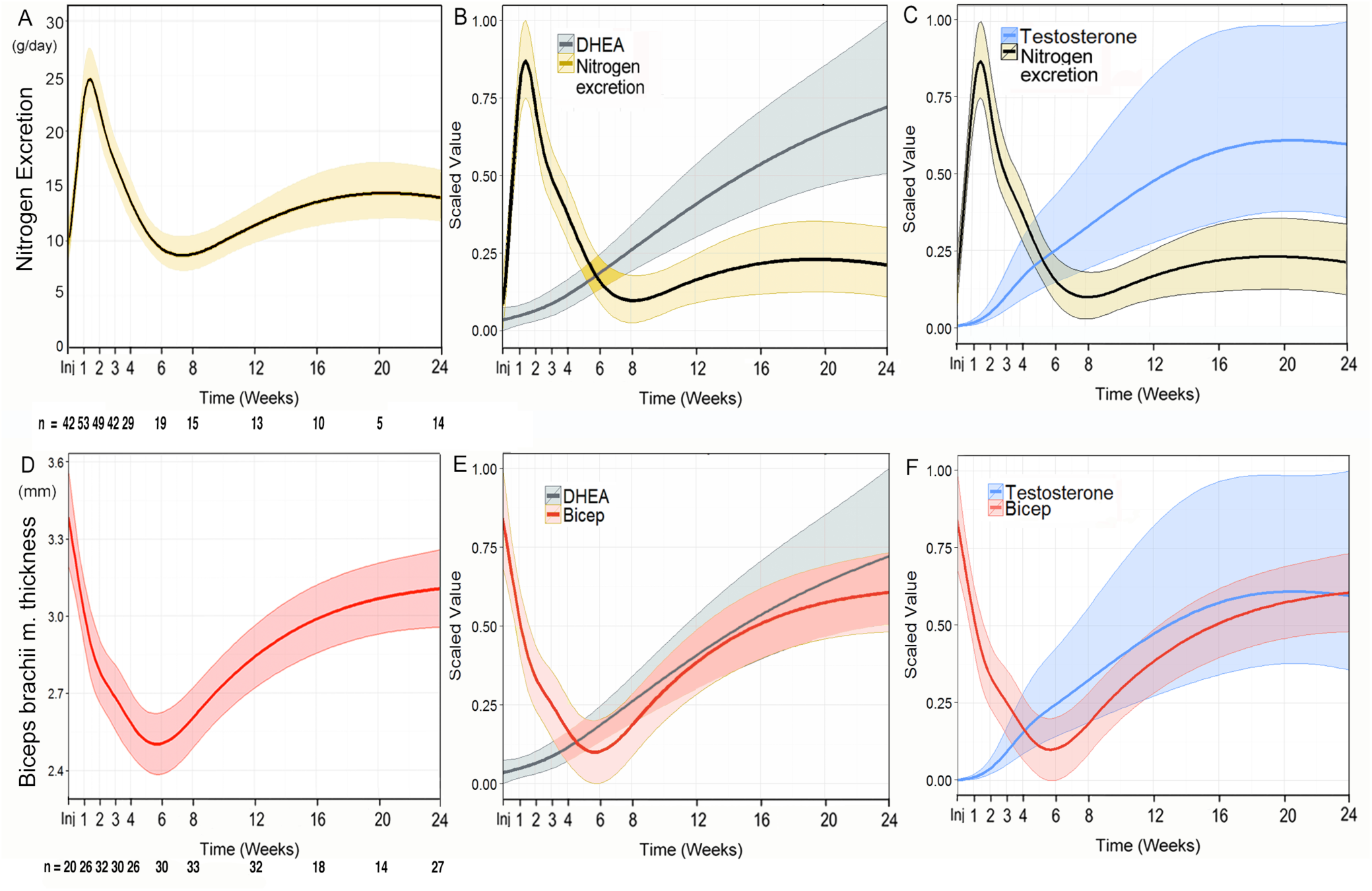
The relationship between (A) Urinary Nitrogen Excretion or (B) biceps muscle thickness with (B and D) DHEA and (C and F) testosterone, over time for young (<50), severely injured (NISS>15) males who had survived and not been given anabolic steroids. Muscle thickness data was modelled using a mixed effects technique; modelling time as a 6 and 7-knot restricted cubic spline respectively provided the best fit. Data are means and 95% confidence intervals for model-based predicted fixed effects of time are shown.

The biceps brachii muscle was the most reliable site for ultrasound measurement of muscle thickness; dressings, amputations and other wounds hampered the measurements of the other muscle areas. Changes in biceps brachii muscle thickness followed a U-shaped curve after injury, reaching a nadir at 6 weeks (day-1 after trauma compared with week-6, p=0·001). The mean muscle loss was 22.7±12.5% (Fig. 4D). Similar to TUN, muscle thickness recovered alongside gradually increasing adrenal and gonadal androgen production (Fig. 4E+F).

### Clinical Course of Post-Traumatic Recovery and Serum Androgen Dynamics

The relationship between adrenal and gonadal androgens and the Sequential Organ Failure Score (SOFA) and probability of sepsis are illustrated in Fig. 5. During the first four weeks, serum DHEA, DHEAS, and testosterone all correlated with the clinical SOFA score (autocorrelation factor (ACF) = 0·85, 0·90 and −0·79, respectively). The serum concentrations of all three steroids also showed strong associations with the probability of sepsis (R=-0·85, 0·85 and −0·97 for serum DHEA, DHEAS and testosterone, respectively). SOFA score and probability of sepsis also correlated strongly with the DHEA:DHEAS ratio (autocorrelation factor (ACF) = −0·94 and −0·96 respectively) and with the serum cortisol-to-DHEAS ratio, negatively for the SOFA score but positively for probability of sepsis (autocorrelation factor (ACF) = −0·81 and 0·89, respectively) (**Suppl. Fig. 5**) (34).

**Figure 5.**
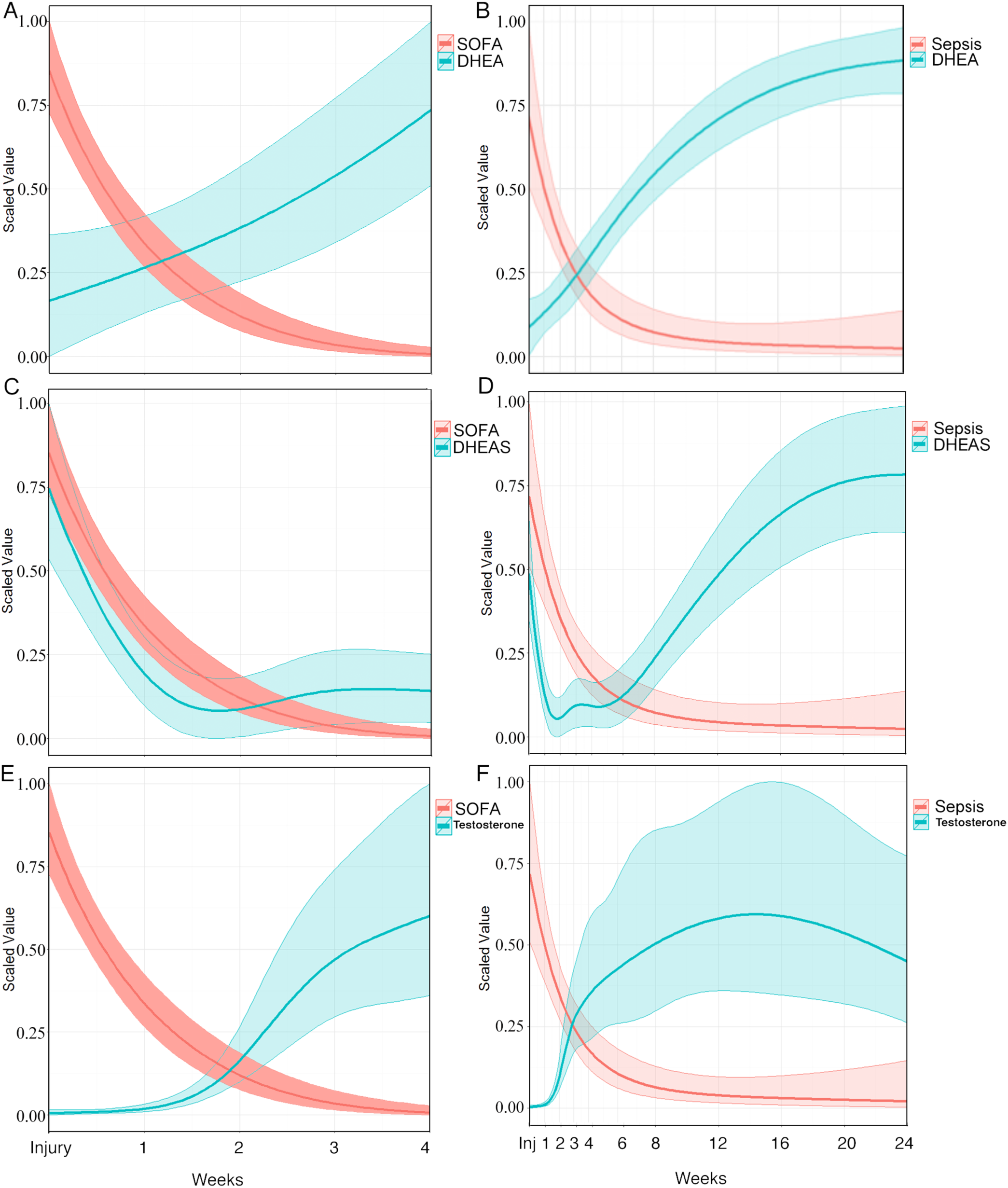
Sequential Organ Failure Assessment (SOFA) score and probability of sepsis in relation to endocrine response. SOFA and sepsis are related serum concentrations of DHEA (Panels A+B), DHEAS (Panels C+D), and testosterone (E+F). Data were modelled using a non-linear mixed effects model that accounts for unbalanced repeated measures using a 4-knot cubic spline. Modelled data are reported as means and 95% confidence intervals.

### Opioid administration and endocrine recovery

To examine whether opioid administration affected endocrine recovery, we modelled the impact of the total cumulative in-patient opioid dose on circulating steroid concentrations during recovery from major trauma. For this purpose, we categorised patients according to cumulative opioid dose. Modelling took into account the differences in ISS, length-of-stay (LOS), ICU LOS, and SOFA score.

The adjusted modelling revealed a dose-dependent impact of opioid treatment, with a higher initial peak of serum cortisol and the cortisol/cortisone ratio in those on higher doses (≥3000mg) while those on lower doses had initially lower serum cortisol concentrations but showed better recovery of cortisol and cortisol/cortisone 2 months into the recovery period, with broad interindividual variability in those with high cumulative opioid doses (Fig. 6).

**Figure 6.**
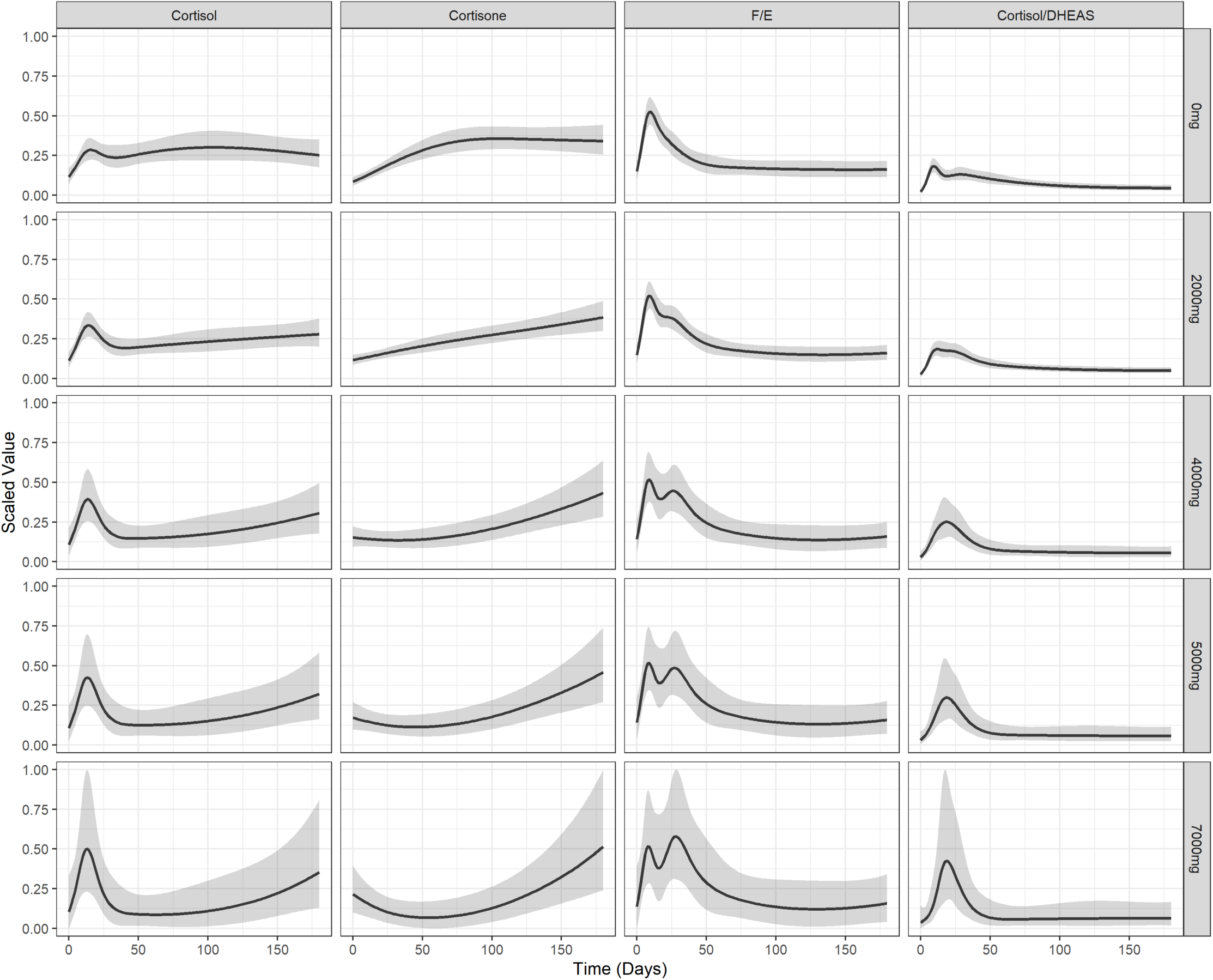
Impact of total inpatient opioid dose on circulating glucocorticoids after major trauma. Serum concentrations are scaled for cortisol, cortisone, the cortisol-to-cortisone ratio and the cortisol-to-DHEAS ratio. Data are represented after modelling of the raw data using a non-linear mixed effects model that accounts for unbalanced repeated measures using a 4-knot cubic spline. Modelled data are shown as means and 95% confidence intervals.

Opioid administration showed a pronounced, dose-dependent effect on adrenal and gonadal androgen production, with significantly delayed recovery of serum DHEA and DHEAS in patients on higher opioid doses (p=0.029, p=<0.001 respectively; Fig. 7). By contrast, serum testosterone concentrations, which were initially equally suppressed in all cumulative dose groups, showed a much faster recovery in individuals who received higher (≥3000mg) total cumulative opioid doses. However, these confidence intervals were large for these model estimates (Fig. 7).

**Figure 7.**
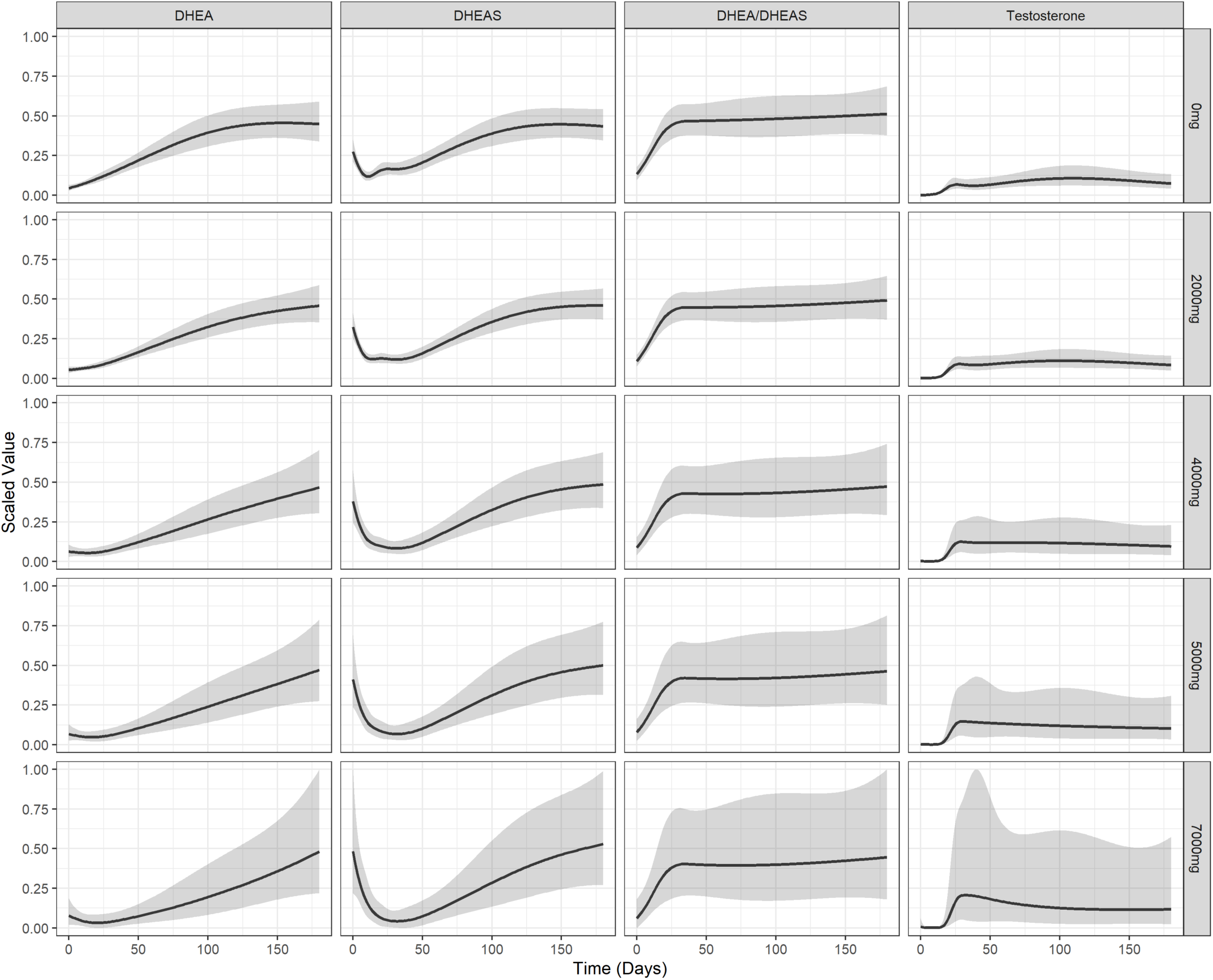
Impact of total inpatient opioid dose on serum androgen and androgen precursors after major trauma. Serum concentrations are scaled for DHEA, DHEAS, the DHEA/ DHEAS ratio, and testosterone. Data are represented after modelling of the raw data using a non-linear mixed effects model that accounts for unbalanced repeated measures using a 4-knot cubic spline. Modelled data are shown as means and 95% confidence intervals.

## DISCUSSION

In this study, we have characterized the response of adrenal and gonadal steroids and catabolic metabolism to severe injury, describing the related dynamic changes for six months post-injury. Modelling the data has allowed us to provide a detailed description of the transition from catabolism to anabolism during recovery from severe injury, including investigating the impact of cumulative in-patient opioid dose. Our data are the first to provide detailed adrenal and gonadal steroids beyond the first days after trauma in a large cohort of young patients, with all patients recruited prospectively and steroid analysis carried out by tandem mass spectrometry.

As summarized in a recent meta-analysis (35), previous data on serum cortisol after injury are limited to small cohorts derived from elective surgery, rarely followed up for more than two days. In our study, serum cortisol quickly returned to normal following slight initial increases after acute trauma. In contrast, serum cortisone remained low for three months post-injury. Our study revealed an initial phase of minor glucocorticoid activation with a transient increase in the serum cortisol-to-cortisone ratio, with changes in urinary glucocorticoid metabolites indicative of increased 11β-HSD1 activity. The cortisol-activating enzyme 11β-HSD1 is the major enzyme converting inactive cortisone to cortisol and has been shown to be upregulated systemically and locally in response to inflammation, thereby dampening the inflammatory response (36, 37). Skeletal muscle expresses 11β-HSD1 (38), Previous studies reported increased 11β-HSD1 activity in an animal model of trauma haemorrhage (39). and improved wound healing in mice treated with 11β-HSD1 inhibitors (40). However, human data after trauma are lacking. There is substantial evidence indicating a reduced cortisol clearance in critical illness, due to decreased cortisol inactivation in liver and kidney (41), this mechanism could also be responsible for the slight changes in cortisol and cortisone we observed. This was corroborated by the observed reduction in the urinary cortisol-to-cortisone ratio, which is reflective of 11β-HSD2 activity.

Interestingly, patients on higher opioid doses, showed a higher early peak in cortisol production after trauma, followed by persistently lower circulating cortisol during the recovery period, as compared with patients on lower opioid doses. Previous reports have described suppressive effects of opioids on the HPA axis, though studies in smaller mammals have indicated an acute stimulatory effect of opioid administration on serum cortisol concentrations (42, 43).

We observed a pronounced and sustained loss of adrenal and gonadal androgen synthesis within the first 24 hours following acute major trauma. The recovery of circulating DHEA and testosterone concentrations took two and four months post-injury, respectively, and DHEAS remained pathologically suppressed at the end of the six-month follow-up period. In a mouse model of acute inflammation, sustained suppression of the expression of the DHEA sulfotransferase SULT2A1 and its sulfate donor enzyme, PAPSS2, have been described (44). We reviewed 23 previous studies that measured serum DHEA and DHEAS in critically ill patients (**Suppl. Table 1**) (45), but most studies followed patients for only a few days and relatively few patients suffered from acute trauma. One previous study measured both serum DHEA and DHEAS in 181 patients with septic shock, and 31 patients with acute hip fracture (46). Serum DHEAS was decreased in both groups, while DHEA was increased in sepsis but decreased after trauma. This suggested an inflammation-mediated downregulation of DHEA sulfation after trauma, resulting in a dissociation of serum DHEA and DHEAS. In our study, this was also observed, as indicated by a sustained increase in the serum DHEA/DHEAS ratio and persistently low serum DHEAS concentrations.

A number of previous studies have described an association of infection and mortality with low circulating DHEAS concentrations and a raised serum cortisol-to-DHEAS ratio in patients with trauma (47–50). In vitro studies have demonstrated that cortisol decreases neutrophil superoxide production, which is counteracted by coincubation with DHEAS (47). Furthermore, we have previously shown that DHEAS, but not DHEA, directly enhances neutrophil superoxide generation; a key mechanism of human bactericidal function via activation of protein kinase C-β, independent of androgen receptor signalling (51). In the present study, carried out in severely injured men younger than 50 years of age, we observed suppression of both serum DHEA and DHEAS post-injury, indicating that the loss of adrenal androgen synthesis is a trauma-related event. Importantly, we showed for the first time that this decrease in circulating adrenal androgen precursors is sustained for several months, and that DHEAS remains low even six months post-injury.

Alongside the decrease in adrenal androgen synthesis, we observed a near complete loss of gonadal testosterone production and pituitary LH secretion immediately after trauma. Both the gradual recovery of adrenal and gonadal androgen production paralleled the decrease in catabolism, as assessed by urinary nitrogen excretion and biceps muscle thickness. The suppression of the hypothalamus-pituitary-gonadal (HPG) axis after severe injury shown in our study is supported by the literature (52–55). Our prospective, longitudinal data demonstrate that suppression of the HPG axis is of shorter duration than that of the HPA axis. In traumatic brain injury studies, a significant proportion of patients go on to develop anterior pituitary dysfunction including secondary hypogonadism (56). However, in our study traumatic brain injury was an exclusion criterion. While limited data from patients with burns and critical illness have suggested a central, hypothalamic-pituitary cause of trauma-related hypogonadism (57, 58), the evidence prior to our study has been limited. Our data indicated a central cause of suppression to the gonadotrophic axis, with a decrease in both pituitary LH and gonadal testosterone. Interestingly, we observed a differential impact of the cumulative opioid dose on adrenal and gonadal androgens, respectively, with a significantly delayed recovery of DHEA and DHEAS, but a trend towards faster recovery of gonadal testosterone synthesis in patients with higher cumulative opioid doses. Previous data on opioid effects on adrenal androgen production are very scarce, but our findings with respect to gonadal testosterone biosynthesis contrasted previous studies describing suppressive opioid effects on the HPG axis (42, 43).

Our study revealed a loss of both adrenal and gonadal androgen production in young and middle-aged men after major trauma. This effect was further enhanced by long-lasting suppression of androgen-activating systemic 5a-reductase activity, as demonstrated by urinary steroid metabolite analysis. Androgens are important in wound healing, erythrocytosis, bone density and muscle mass (59). The catabolic state that occurs following trauma thus presents a significant challenge. The use of androgens to ameliorate catabolism has some precedent, as evidenced by the use of the synthetic androgen, oxandrolone, that has some proven benefit in treating burn injury (10). A meta-analysis of 15 Randomised Controlled Trials including 806 burns patients by Li et al, showed significant benefits (P < 0.05) for using oxandrolone, including less net weight loss, lean body mass loss, nitrogen loss, donor-site healing time, and length of stay in the catabolic and rehabilitative phases (60). The use of oxandrolone in major trauma was investigated in two intensive care studies but no benefit was demonstrated (61, 62).

The strengths of our study include its prospective nature, narrow age range of the patients, single gender, single site for recruitment and analysis and detailed follow-up over six months as well as the measurement of circulating (and in a smaller cohort also excreted) steroid hormones by state-of-the-art mass spectrometry assays. Analysing a young to middle-aged patient cohort has also reduced the confounding effects of age-related co-morbidities. Another strength is the unique opportunity our study offered for analysis of the opioid effects on endocrine recovery, facilitated by detailed prospective, longitudinal phenotyping with dedicated software.

Our study was limited by the diverse nature of major trauma patients in relation to injury pattern and the involvement of military casualties. The timing and number of observations during our study was pragmatic and some statistical comparisons were made using modelled data. While we measured total cortisol and cortisone by tandem mass spectrometry, we did not measure free cortisol or cortisol-binding globulin. We were only able to measure urinary steroid excretion in a sub-cohort of patients, as accurate and repeated collection of 24-h urine proved very challenging under ICU conditions. The estimation of nitrogen excretion was pragmatic due to the diverse nature of the patients and we were not able to record nitrogen intake. Ultrasound estimation of muscle thickness was performed at four different body sites, but many individuals had limbs missing or extensive wounds that prevented measurements. While imperfect, the longitudinal nature of these measurements allowed us to model these changes over time.

In conclusion, in this most detailed and first prospective study of the steroid response to major trauma, we followed the patients from severe injury to six months of recovery, revealing pronounced and sustained decreases in adrenal and gonadal androgen biosynthesis. Recovery of androgen production in the severely injured patients was mirrored by a switch from catabolism to anabolism as reflected by recovery of muscle mass and a decrease in nitrogen loss. Adrenal and gonadal androgens correlated with risk of sepsis. It is tempting to suggest that an anabolic intervention with androgens or androgen precursors could have a beneficial effect on health outcomes during recovery from major trauma. However, this will need to be investigated by future intervention studies.

## Acknowledgements

The SIR Study was part of the Surgeon General’s Casualty Nutrition Study (SGCNS), supported by University Hospitals Birmingham NHS Foundation Trust (UHB) and the University of Birmingham. Additional funding was provided by the Drummond Trust Foundation. WA and JM received support from the National Institute for Health Research (NIHR) Birmingham Biomedical Research Centre (Grant Reference Number BRC-1215-20009). The views expressed are those of the authors not necessarily those of the NIHR or the Department of Health and Social Care UK.

We thank staff from the NIHR Surgical Reconstruction and Microbiology Centre, Leah Duffy, Lauren Cooper, Peter Ip and Aisling Crombie; military nurses Michelle Taylor and Matt Daley; junior doctors from UHB, Abigail Routledge, Ben Booth, Rob Staruch and Researchers and staff from University of Birmingham, Hema Chahal, Donna M. O’Neil, Jon Hazeldine and Pete Hampson; Institute of Naval Medicine, Dr Adrian Allsopp, Sophie Britland, Dr Pieter Brown, Roz Cobley, Simon Delves, Anneliese Shaw, Dr Fran Gunner, Jenny Hayward-Karlsson; Defence Rehabilitation Centre Headley Court, Jakob Kristensen, Wg Cdr Alex Bennett; University of Surrey, Prof Susan Lanham-New; Imperial University, Prof Stephen Brett; Dr Kevin Murphy, Prof Gary Frost; members of SGCNS not already mentioned, Surg Cdr Jane Risdall, Andy Roberts, Lt Col Sandra Williams, and Col Duncan Wilson.

---

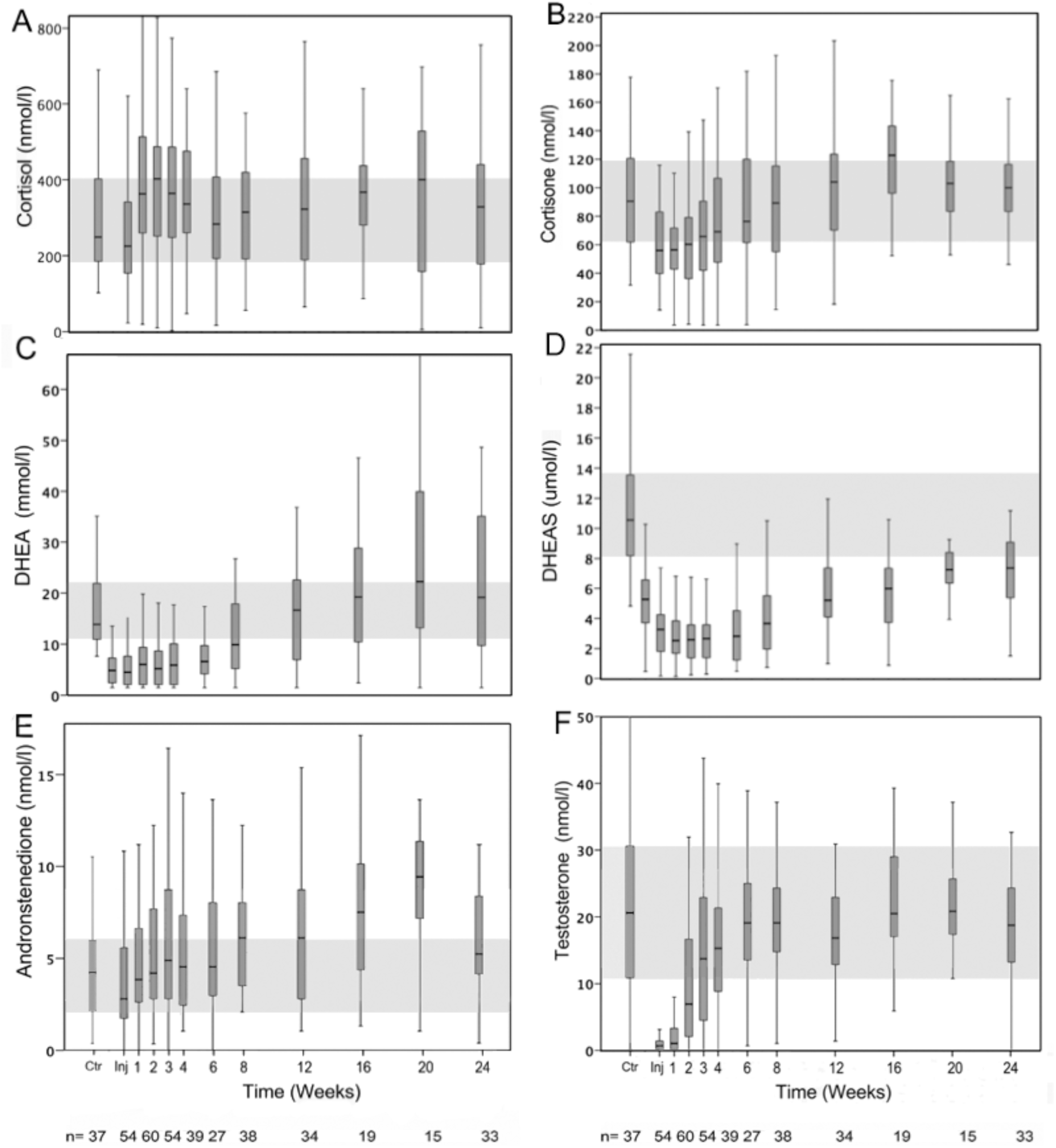

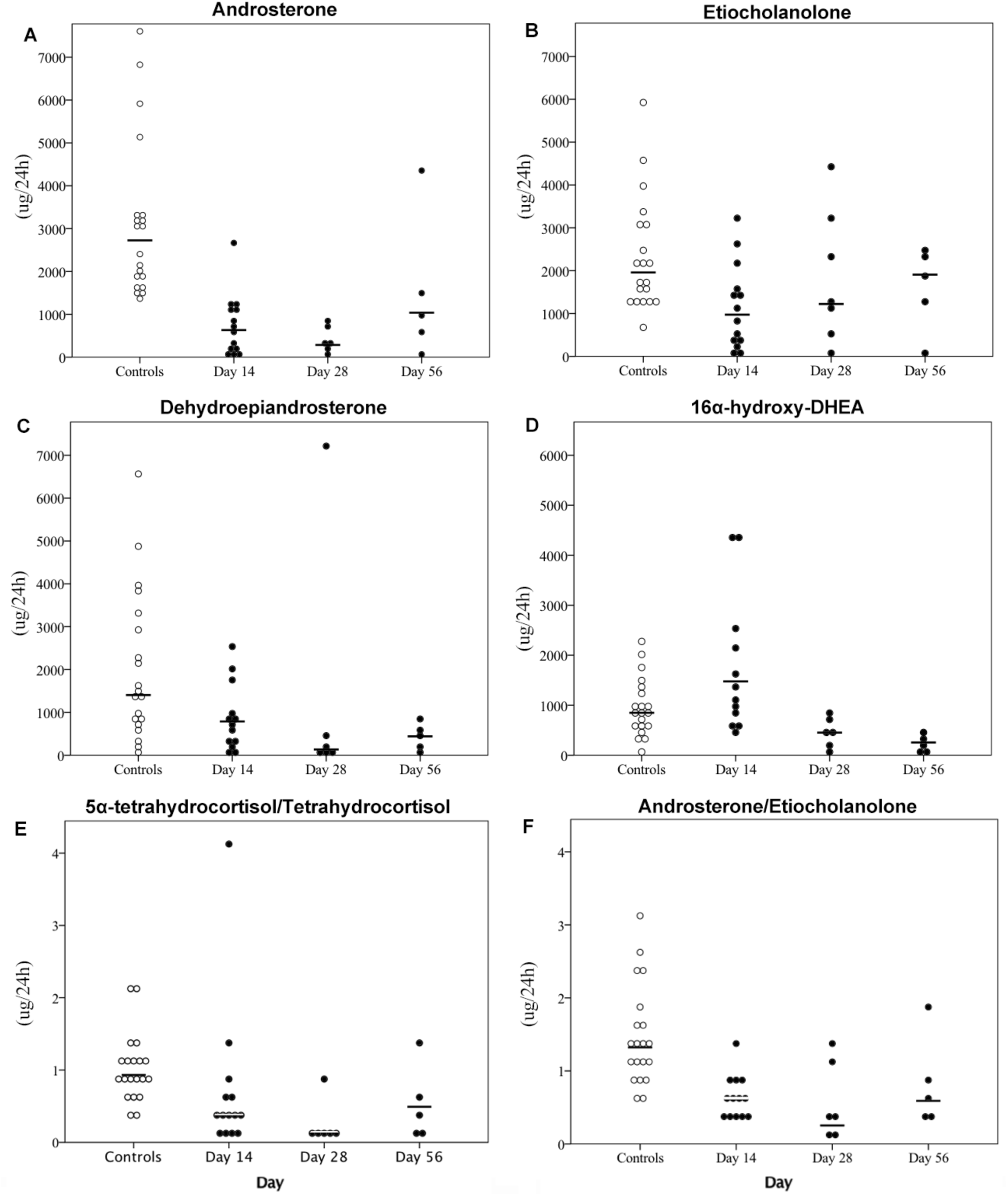

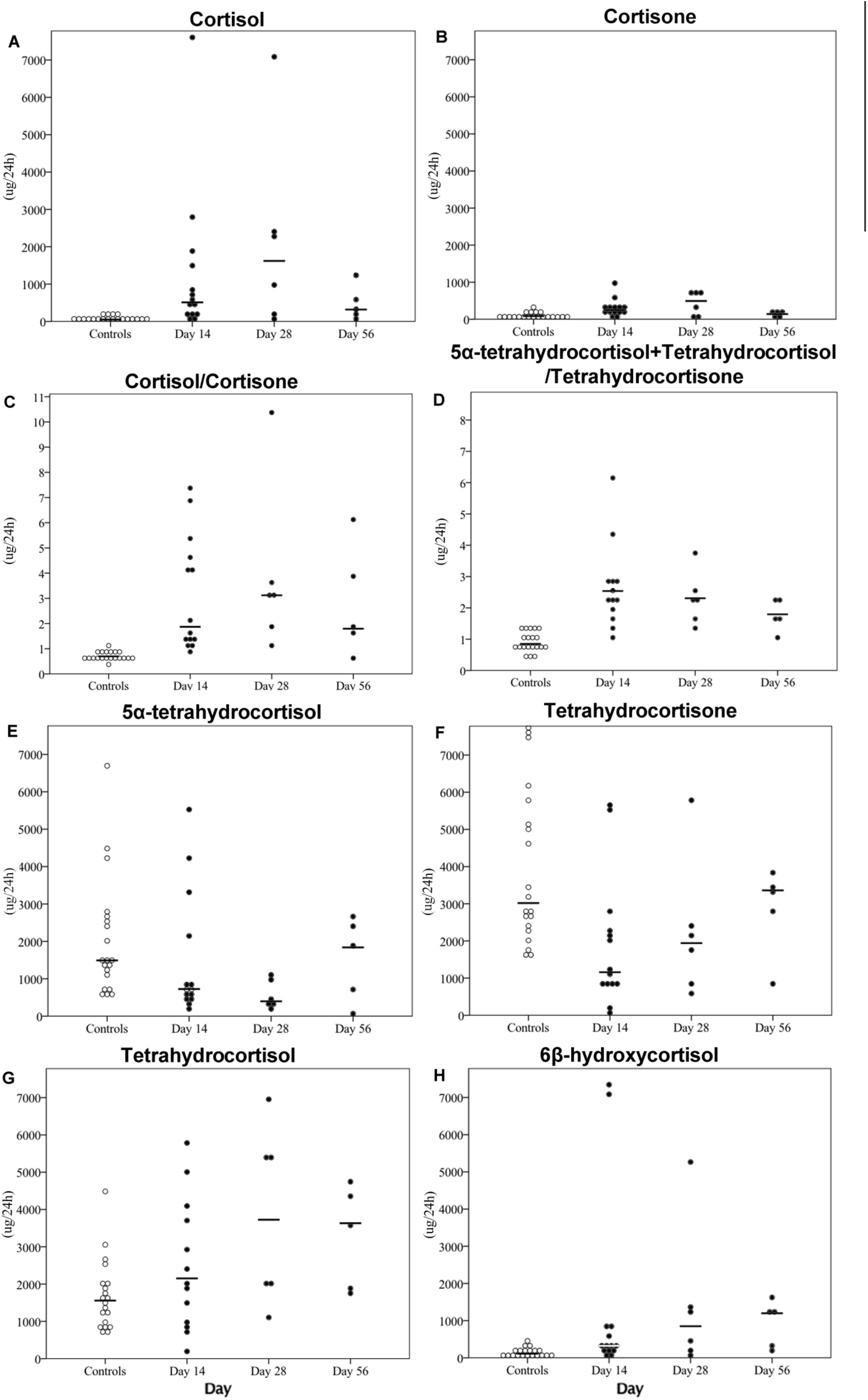

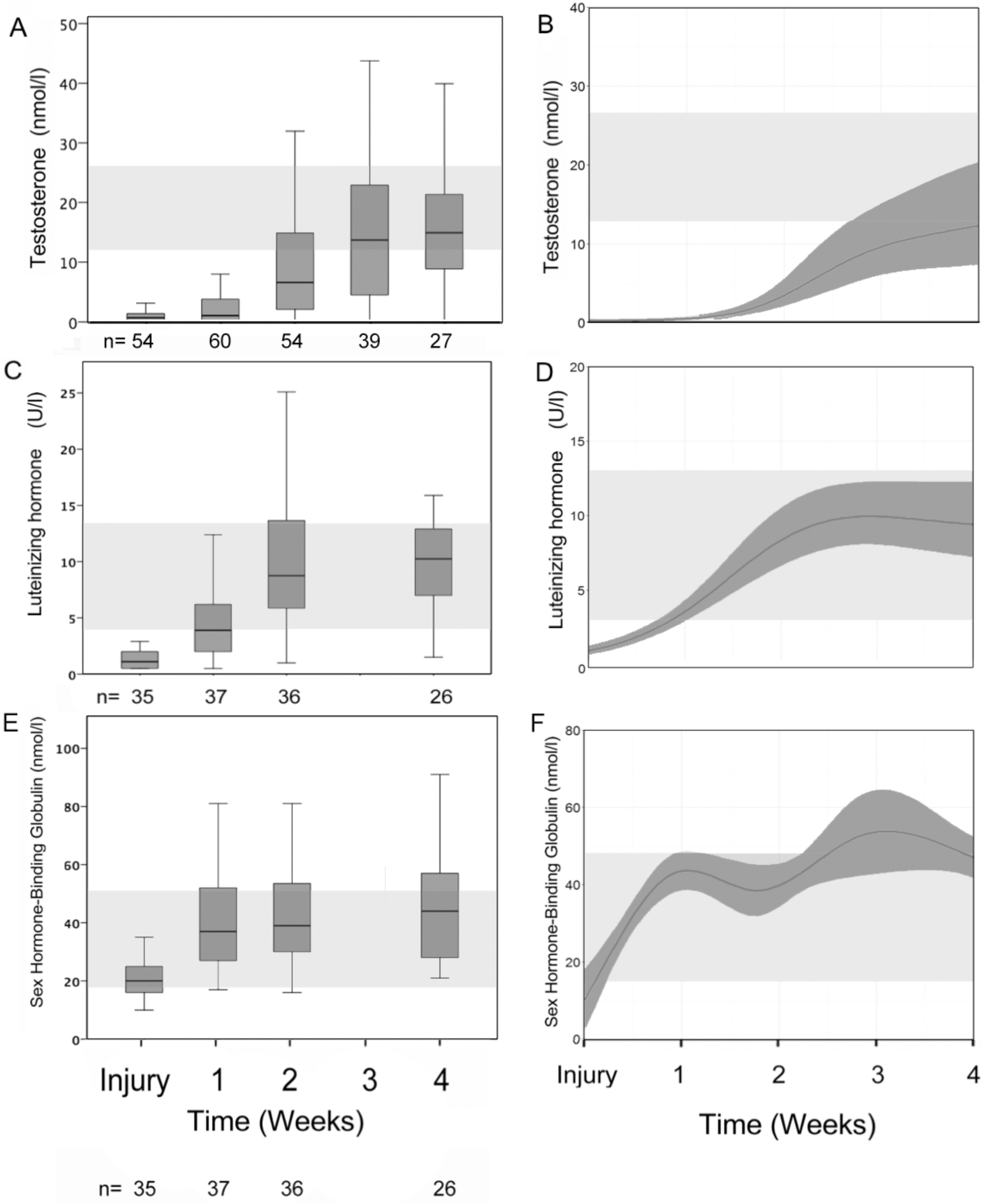

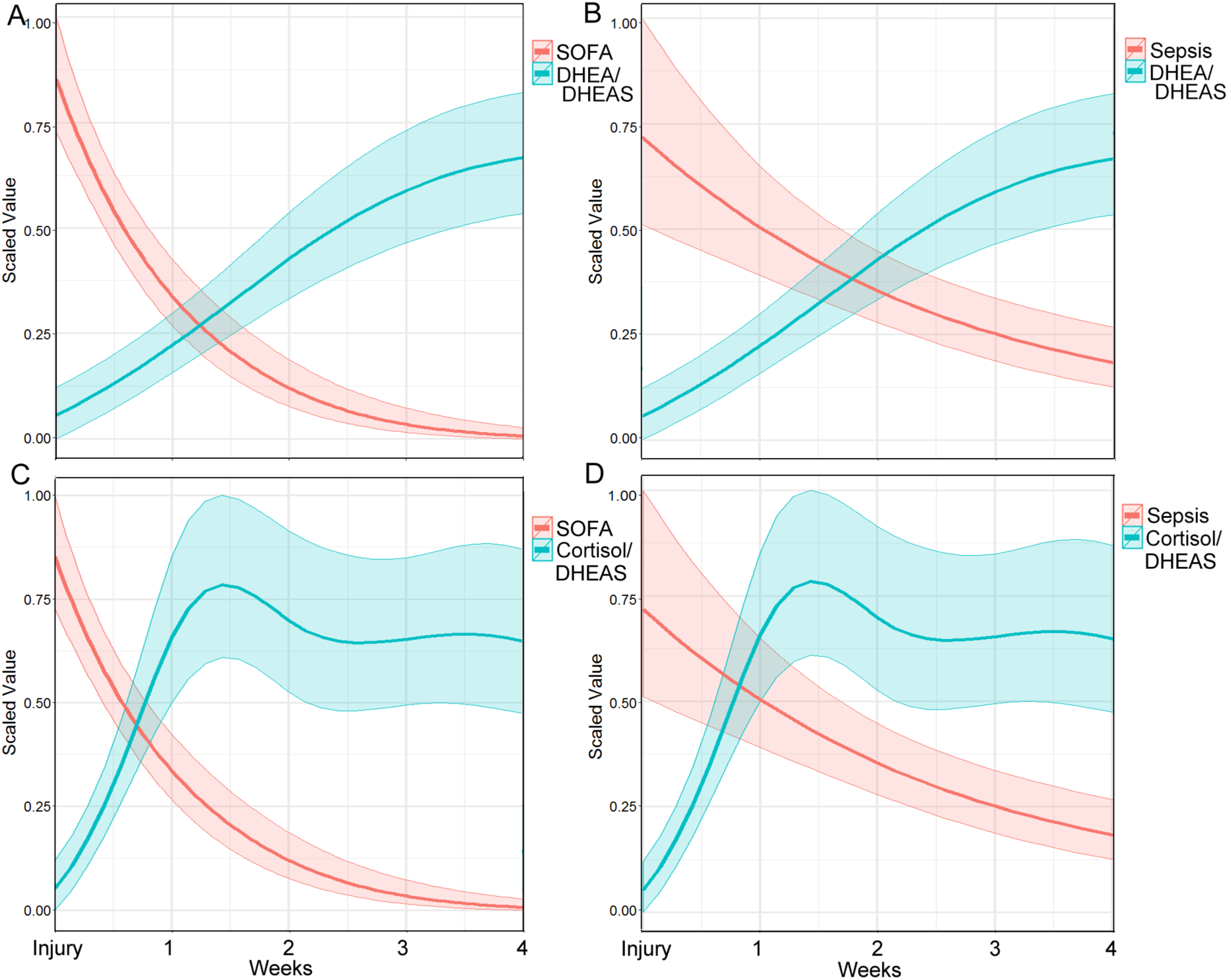

**Suppl. Table 1.** Longitudinal Studies in critically ill patients analysing DHEA and DHEAS.

| Author | Year | Population | Measurements | Comparison | Outcome |
| --- | --- | --- | --- | --- | --- |
| Kolditz (38) | 2010 | 59 patients hospitalized pneumonia | CRP, IL6, TNFa, cortisol baseline and after simulation, ACTH, DHEA, DHEAS on the morning of hospital admission | Clinically stable on day 4 vs clinically unstable on day 4 | DHEA/DHEAS ratio was significantly different – higher in stable patients (ratio of 11 vs 7) |
| Sharshar (39) | 2010 | 103 ventilated ICU patients | IGF, prolactin, TSH, FSH ,LH, estradiol, testosterone, DHEA, DHEAS, cortisol on day 1 when 7 days after wakening | Correlation to SAPSII F vs M | SAPS II assessed at awakening was inversely correlated with plasma levels of DHEA |
| Klouche(40) | 2007 | 36 ICU patients with critical illness, severe sepsis or septic shock | 24h after admission, samples at 0800hrs for cortisol, ACTH, and DHEAS | Elderly vs nonelderly | DHEAS 0.52 nmol/l vs 0.93 in >75 year old vs <75 years old |
| Chinga-Alayo(41) | 2005 | 113 ICU patients | First hour after admission to ICU – cortisol, thyrotropin, t3, t4, cortisol, prolactin, GH, DHEA | 88 Survivors vs 25 non-survivors | No difference in DHEA ( 86 vs 73 µg/dl) |
| Butcher(42) | 2005 | 35 elderly and 9 young patients with hip fracture undergoing surgery | serum adrenal stress hormones DHEAS and cortisol immediately following injury | Young vs elderly | DHEAS was higher young trauma group ($6.29 \pm 2.41 \mu M$) compared with elderly hip fracture patients ($1.82 \pm 1.58 \mu M$). |
| Beishuizen(43) | 2002 | 30 patients with septic shock, 8 with multiple trauma, and 40 controls | serial measurements of DHEAS , cortisol, TNFa, IL-6, and ACTH over 14 days or until death. | Septic shock vs multiple trauma vs controls, Survivors vs non-survivors | DHEAS was lower in septic shock than trauma, both lower control. DHEAS -ve correlation with age, IL-6 and APACHE II scores in both patient groups. DHEAS lower in non-survivors. |
| Sharshar(44) | 2011 | 102 patients in ICU who regained consciousness after 7 days ventilation | IGF-1, prolactin, TSH, FSH, LH, estradiol, progesterone, test-, DHEA, DHEAS, cortisol 1<sup>st</sup> day awake | Survivors vs non-survivors | DHEA and DHEAS were different in men but not in women (lower in non-survivors). |
| Van den Berghe(45) | 2002 | 33 men long critical illness and age 50, BMI - controls. RCT 5 days of GHRP-2, GHRP2 + TRH infusion or pulsatile | IGF-I, IGFbPs, thyroid hormones, gonadal and adrenal steroids, proinflammatory, metabolic and inflammation markers daily. | Comparison between different therapies | Neither of the foregoing adrenal steroids was altered by any of the study drugs. Replacement requires replacement of somatotrophic, thyrotrophic and gonadotrophic axes to reduce catabolism. |
| Spratt(46) | 1993 | postmenopausal women; 20 with acute critical illness vs 110 healthy controls | Day 1-5 DHEA, DHEAS, androstenedione, testosterone, estrogen, gonadotropins, cortisol | Patients vs controls | Admission levels of DHEA and DHEAS were not elevated in patients. Serum DHEA decreased by day 5 in with gonadotropins |
| Gottschlich(47) | 2009 | 40 patients age 3–18 %TBSA 50.1 were randomly assigned to zolpidem or haloperidol. | 2 week study; epinephrine, norepi-, GH, melatonin, DHEA, serotonin, cortisol -0600hrs each study day. | Therapy 1 vs 2 | Both drugs were associated with increased DHEA levels ($P < .03$); no other hormones were affected by medication . |
| Arlt (48) | 2006 | cross-sectional study consisting of 181 patients with septic shock, 31 patients with acute trauma, and 60 healthy controls. | cortisol, DHEA, and DHEAS were measured before and 60 min after ACTH stimulation. | Septic shock vs acute trauma vs healthy controls | DHEAS lower in septic shock & trauma compared to controls, DHEA was increased in sepsis, decreased in trauma. Sepsis; cortisol and DHEA not increased after ACTH. Most severely ill higher cortisol:DHEA. Cortisol:DHEA increased non-survivors septic shock. |
| Mueller (49) | 2014 | 179 patients hospitalized with community acquired pneumonia. | DHEA, DHEAS, cortisol | Correlation to PSI score | Correlation between PSI score and DHEAS, cortisol/DHEA, cortisol/DHEAS and DHEA/DHEAS. In age, gender adjusted analysis, DHEA but not DHEAS associated with all-cause mortality. |
| Ven den Berghe(50) | 1995 | 20 critically ill polytrauma receiving dopamine were studied to evaluate dopamine withdrawal | DHEAS, prolactin, cortisol during dopamine infusion | Dopamine withdrawal vs continued dopamine | Withdrawal of dopamine 25% increase DHEAS within 24hrs, not DHEAS levels when dopa- continued 2x nights. Prolactin undetectable dopa- infused, increased after 24 hrs of withdrawal. |
| Dimopoulou (51) | 2007 | 203 severely ill -trauma(93), medical(57), or surgical (53). | 24h of admission ICU AM sample to measure cortisol, ACTH and DHEAS. | 149 survivors and 54 nonsurvivors | Nonsurvivors had a lower incremental rise in DHEAS (1065 vs. 1642 ng/ml) than survivors. |
| Osorio(52) | 2002 | 38 patients scheduled for cholecystectomy | DHEAS, ACTH, cortisol, hGH, IGF-1, and IGFBP-3 preop, and then 2 and 7 days after surgery. | Preop to day 2 to day 7 after surgery | Reduction in DHEAS on days 2 and 7 after surgery versus the preop values . |
| Almoosa(53) | 2014 | 30 men ventilated for >24 hrs for acute respiratory failure. | Blood samples on ICU day 1 and day 3, serum testosterone. | Day 1 and 3 levels, compared to reference | DHEAS levels normal. Total and free testosterone correlated inversely ventilator days and ICU LOS. |
| Dolecek (54) | 2006 | 29 polytraumas and 28 burned patients (evaluated by their Burn Index, BI) were followed | Bone markers , iPTH, calcium, inorg PO4- , 25OH vit. D3, testosterone, DHT, free test- , cortisol, 17β estradiol, DHEAS, TNFα, cytokines. | days 1-7-14-28, of the burned at 1-7-14-28-56, as well as after 6 and 12 months. | All androgens (T, DHT, FT) decreased significantly in the males, DHEAS decreased in male and females. |
| Dossett(55) | 2008 | 991 injured patients remaining in the ICU for at least 48 hours | Sex hormones (estradiol, progesterone, testosterone, prolactin, and DHEAS) | Survivors vs non-survivors | Estradiol elevated in nonsurvivors . Estradiol most severely injured had highest. Progesterone, test-, DHEA-S were higher in nonsurvivors; ability to accurately predict death lower than estradiol . |
| Folan(56) | 2001 | 191 men and women in ICU | DHEA, DHEA-S, and cortisol within 24 hrs of admission and compared with admission APACHE II scores. | APACHE 2 scores and Surgical(SICU) and Medical ICU | The correlations between APACHE II scores and DHEA data for women in the ICU correlation between APACHE II and DHEAS women in SICU. |
| Ilias (57) | 2007 | 83 men and 11 women with multiple trauma | TSH, fT4, T3, ACTH, prolactin, cortisol and DHEAS. | Survivors vs non-survivors | ACTH and DHEAS higher survivors. APACHE II and Cortisol post-Synacthen, DHEAS, TSH*age assessed survival/non-survival better than APACHE II, SOFA or IS scores alone. |
| Brorsson(58) | 2014 | 50 trauma patients admitted to a level-1-trauma centre | Serum & saliva cortisol from injury to five days after trauma. CBG, DHEA & DHEAS twice 5 days after trauma. | Temporal change | A significant decrease over time was observed in DHEA, and DHEAS. |
| Bergquist (59) | 2016 | 16 adult male patients with burn injury (14.5–72%TBSA ) | Plasma cortisol, cortisone, corticosterone, 11-deoxycortisol, DHEA, androstenedione, test-, pregnenolone, , estrone and estradiol and progesterone. | Changes 0, 1, 3, 7, 14, 21 days post burn | Burn injury alters endogenous steroid biosynthesis, reduce testosterone and DHEA levels to 3 weeks and elevated estrone concentrations post injury. Concentrations of glucocorticoids, progestagens and androgen precursors correlated positively TBSA. |

